# *90-90-90, epidemic control* and *Ending AIDS*: global situation and recommendations

**DOI:** 10.1101/196972

**Authors:** Reuben Granich, Somya Gupta, Brian Williams

## HIV remains a devastating pandemic

Although human immunodeficiency virus (HIV) first came to our attention thirty-seven years ago, the acquired immunodeficiency syndrome (AIDS) which it causes is, without treatment, 100% fatal with devastating consequences for millions of people^1^. More than 35 million people have already died of AIDS-related conditions and HIV/AIDS is ranked the third largest pandemic after the 14^th^ century *Black Death* and the 1918 influenza pandemic. HIV continues to spread and an estimated 37 million people are living with HIV (PLHIV) globally^2^. Ending the pandemic of HIV and acquired immunodeficiency syndrome (AIDS) is necessary and feasible but poses important challenges.

By 2016, an unprecedented global response has provided anti-retroviral therapy (ART) for 19.5 (53%) million people living with HIV. Despite this progress, in 2016, 1.8 million people were newly infected and 1 million people had died of AIDS-related conditions^2,3,4^. Additionally, stigma and discrimination lead to social neglect, putting vulnerable and marginalized people at risk and compromising their ability to access HIV services. HIV infection is the strongest risk factor for tuberculosis (TB) resulting in an estimated 1.2 million HIV-associated TB cases and 400 thousand HIV-associated TB deaths in 2015^5^ even though HIV-associated TB is almost entirely preventable through treatment for HIV and curable if properly treated^6^. In addition to preventing illness, people who are on ART and virally suppressed do not transmit the virus^7^. Providing ART to everyone infected with HIV could avert the majority of illnesses, deaths and infections, preventing suffering for millions of individuals, their partners, and the community.

In this article we examine the concepts of 90-90-90, epidemic control and ending AIDS. We review the global situation including innovations that will likely have a major impact and make recommendations regarding the global HIV response. Although there has been considerable debate on the topic of controlling HIV and ending AIDS, very few articles have examined the issue in depth from both an epidemiologic and political perspective. Specifically, this paper reviews the epidemiologic criteria for ending AIDS while placing it in the political context of the global HIV response. We provide a framework for understanding the history of the HIV response while addressing the issue of treatment as prevention, economics of ending AIDS, major innovations, and last mile issues.

## Ending AIDS

To better understand the concept of *Ending AIDS*, it is helpful to consider the HIV response as being characterized by three overlapping phases (Figure 1): 1) *Devastation* 2) *Discovery* and 3) *Ending AIDS*. After HIV was first discovered in 1981, it spread across the globe with Africa bearing the brunt of the pandemic^1^. The social response was weak with denial being the dominant stance, stigma was widespread, and people were often left to die alone. Families and friends struggled in isolation leaving a few health care providers to deal with the sick and dying while governments and other authorities often ignored or exacerbated the problem^8,9^. The devastation was profound and difficult to capture in words—in many settings everyone lost a loved one or family member, health, care givers lost most of their patients, and the loss of so many young people in the prime of their lives changed society. Hospitals were filled to capacity and, without HIV treatment, it was not uncommon to have one or two deaths a day on tuberculosis and other wards. Although hundreds of millions of people became ill, over 39 million people have died, and the suffering has been immeasurable, the silver lining was the development of an unprecedented individual and community response. People affected by HIV were the first responders who, while caring for the sick and dying, demanded leadership and resources from local, state and national governments ^8,10,11,12^.

**Figure 1:**
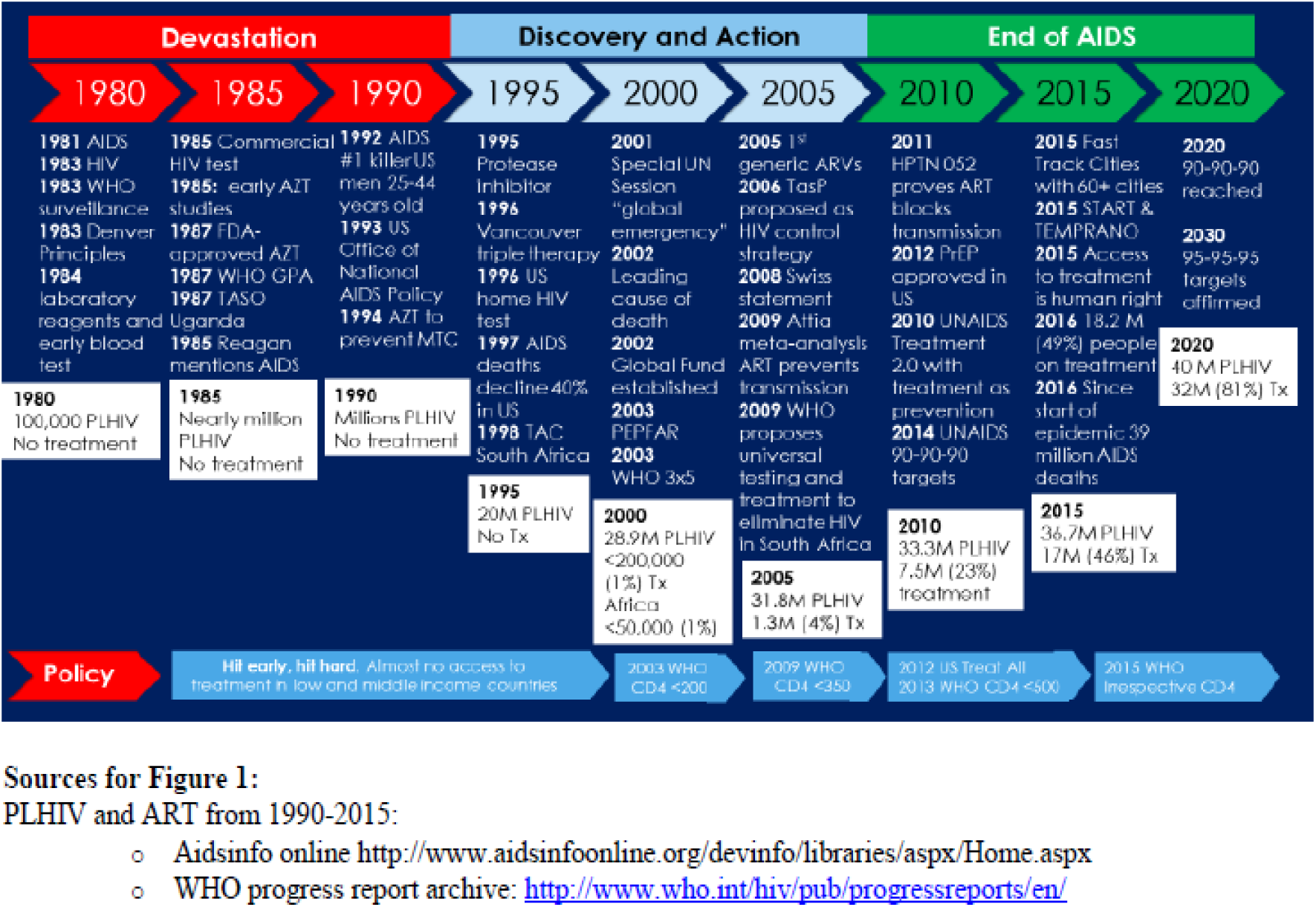
Devastation, Discovery and Action, End of AIDS: Key HIV epidemic milestone from 1980 to 2020

The *Discovery Phase* resulted from the growing collective community, scientific, and political response that unraveled the secrets of HIV more rapidly than any other pathogen in history. Advances included HIV diagnosis, global surveillance systems, improved clinical care and prophylaxis for opportunistic infections, and the discovery of anti-retroviral agents. Community organizations provided care while demanding more resources in the hopes of finding successful treatments, vaccines, or a cure. Nevertheless, the consensus among scientists, policy makers, and community leaders was that ending AIDS was not feasible without a vaccine or cure. The central dogma focused on offering care to help slow HIV progression and ameliorate a painful death, calling for human rights and ending stigma and discrimination, building community support for people with HIV, and trying to control transmission through traditional prevention methods including abstinence, behavior change, syringe exchange and condom use. The development of triple combination therapy for HIV in 1996 represented a significant paradigm shift with the prospect of life-saving treatment that stop the progression of HIV and render them non-infectious to others^13,14^. This was accompanied by demands for increased access to treatment and the realization that it was morally unacceptable to only provide it for wealthy people in economically advantaged countries^10,11^.

By June 2016 an unprecedented international, national and local effort translated the scientific discovery of life-saving treatment into access for 19.5 million people—half of the 37 million people living with HIV^15^. In 2017 this number grew to 22 million people on ART. As access expanded internationally, a number of observational and randomized clinical control studies demonstrated that early antiretroviral therapy (ART) prevents illness, mortality and HIV transmission^7,16-21,22^. Advances included regimens with less frequent dosing, fewer side effects, greater durability and reduced likelihood of drug-resistance. Additionally, epidemiologic modeling using existing scientific data suggested that treatment should be started as soon as possible, irrespective of CD4 cell count, and that expanded access to treatment could lead to HIV control and the eventual elimination of HIV in many settings^23^. Several studies showed that early treatment could provide a near normal life-span and reduce the transmission of HIV to nearly zero^7,20-22^. Increasing treatment coverage in a community was associated with both reduced individual risk and decreased overall community incidence^16,21,24,25^ and some scientists began to call for immediate treatment to save lives and stop the epidemic^18^. Estimations and modeling suggested that front-loading the investment in earlier HIV diagnosis and treatment would be cost-saving over the medium to long term^18,26^. Access to HIV testing was successfully expanded through the use of rapid tests in diverse settings and in multi-disease prevention campaigns and self-testing^27,28,29,30^. The decade brought conclusive evidence that the conventional wisdom embodied by the WHO-recommended *Test-and-Wait* strategy, which called for withholding treatment until the persons immune system was severely compromised, did not make scientific, clinical, public health, human rights or economic sense. These and other developments provided the foundation for the next phase—the *End of AIDS*.

The *End of AIDS* phase builds upon the change in the HIV response from a long term struggle with hopes pinned on a future vaccine or cure to using currently available tools as part of a winnable public health battle. This new phase draws on the observational, community-based studies and modeling that suggest that increasing access to immediate testing and inexpensive treatment reduces illnesses, end AIDS, prevents deaths and eliminates HIV transmission^7,16-25^. It also builds on the success in preventing maternal to child transmission through providing ARVs and more recently treatment through “Option B+” ^31^. A widely accepted definition of HIV elimination is reducing transmission to less than one new infections per thousand population^23^—not to be confused HIV eradication, which would mean the complete removal of HIV from the world, which is not feasible at this time. Even if transmission were stopped completely about 35 million people living with HIV will require treatment for the rest of their natural lives or until a cure or therapeutic vaccine is developed. The shift to ending AIDS also expanded the traditional anti-discrimination and criminalization human rights agenda to include the right to health in the form of HIV treatment^32^.

This emerging consensus around ending AIDS has resulted in new high-level political commitments including United States congressional hearings on “Test and Treat”^33^, an *AIDS free generation^34^*, and a global plan to eliminate mother to child transmission^35^. *PEPFAR 3.0* developed the “right things, right places, right now” strategy to achieve epidemic control through expansion of access to treatment and other interventions in areas with the highest HIV burden^36^. The Joint United Nations Programme on HIV/AIDS (UNAIDS), Presidents Emergency Plan for AIDS Relief (PEPFAR), national governments and cities adopted the *90-90-90* targets: by 2020, 90% of people living with HIV diagnosed, 90% of people diagnosed on sustained ART, and of those, 90% virally suppressed^36,37,38^. These minimal targets translate into 73% of people on treatment and virally suppressed and are designed as a milestone towards the *End of AIDS*. More recently as part of the Sustainable Development Goals, UNAIDS has announced its “Fast Track” commitments and estimates of resource needs ^39^ including increasing resources to 26BN, reducing new infections and deaths to under 500,000, eliminating HIV associated stigma and discrimination and gender inequalities and end all forms of violence and discrimination against women and girls, people living with HIV and key populations by 2020^40^.

The new *End of AIDS* has a dual meaning (Table 1). It represents the abstract political target of ending HIV as a major public health problem but includes achieving the epidemiological target of reducing both AIDS cases and HIV incidence to less than 1 per 1000 population per annum^23, 41^. HIV-associated mortality rates and/or composite mortality and new infection indicators have also been suggested as a measure of epidemiologic control or the *End of AIDS*^36, 42^.

**Table 1:** Definitions for End of AIDS, Epidemiologic control, HIV control, HIV elimination, HIV eradication, and HIV extinction

| Category | Definition | Examples |
| --- | --- | --- |
| End of AIDS (political) | Abstract political target of ending HIV as a major public health problem. Recognition that 35M PLHIV will need support to stay on successful treatment until cure or therapeutic vaccine | Commonly used in political and public health discourse |
| End of AIDS (epidemiological) | Reduction of HIV incidence and AIDS to below one AIDS case per 1000 population<sup>23</sup>.<br>The 90-90-90 and 95-95-95 targets are milestones on the way to the end of AIDS as they translate into 73% and 86% of people being on treatment and virally suppressed, respectively<sup>37</sup>.<br>The Global Plan calls for the elimination of maternal to child transmission to less than 5% transmission<sup>35</sup>. | Leadership in New York State<sup>44</sup>, Cambodia<sup>45</sup>, San Francisco<sup>46</sup>, Vancouver<sup>47</sup> are focused on HIV control, ending AIDS and getting to zero new infections. |
| Epidemiologic Control | The point at which new HIV infections have decreased and fall below the number of AIDS-related deaths | PEPFAR 3.0 |
| HIV control | The reduction of HIV disease incidence, prevalence, morbidity, or mortality to a locally acceptable level as the result of deliberate public health efforts; continued interventions will be needed to maintain the reduction and move towards elimination targets | San Francisco, Vancouver |
| HIV elimination | Reduction of HIV and AIDS in a defined geographical area to below one AIDS case per 1000 people living with HIV per year and a reduction of HIV incidence to 1 new case per 1000 population<sup>23</sup>. Continued intervention measure are required to maintain elimination | Cambodia |
| HIV eradication | Permanent reduction to zero of the worldwide incidence of HIV as a result of deliberate efforts. Intervention measures are no longer needed. | None |
| HIV extinction | The specific agent no longer exists in the laboratory or nature; interventions are no longer needed | None |

\*Table adapted from the Bulletin of the World Health Organization (1988) *The principles of disease elimination and eradication*.<sup>41</sup>

Specifically, measuring the number of deaths per 1000 PLHIV can serve as a reasonable measure of programme effectiveness. Reaching the End of AIDS, as defined by low levels of HIV incidence and AIDS related mortality, has already been achieved in some settings. However, to end AIDS globally will require a near doubling of the number of people on treatment. The new framing is exemplified in the *90-90-90* and *95-95-95* targets as milestones on the way to ending illness or AIDS as they translate into 73% and 86% of people being on treatment and virally suppressed, respectively^37, 43^. The Fast Track Cities initiative embodies the new framing both from the abstract and epidemiologic perspective; over 65 cities are now committed to achieving *90-90-90* and zero stigma^38^. The Global Plan calls for the elimination of mother-to-child transmission^35^. Leadership in New York State^44^, Cambodia^45^, San Francisco^46^, Vancouver^47^ and other settings have used the new science and framing to focus on ending AIDS and getting to zero new infections.

The new *End of AIDS* framing provides an achievable end-game strategy, making a strong business and moral case for increased focus and investment. As part of ending AIDS, earlier access to treatment prevents illness, death and transmission and can reduce costs to the health sector, society, individual patients and the community. In late 2015, following the example of the United States, France, Brazil and seven other countries^48^, WHO recommended starting ART irrespective of CD4 cell count^49^. As of January 2018, 62 countries representing 86% of the global HIV burden have published updated guidelines, reflecting a removal of restrictions or democratization of access to treatment for everyone living with HIV^48^. The next policy challenge necessary to achieving *90-90-90* will be to ensure access to rapid HIV diagnosis through community-based approaches including self-testing^50,51^. Already, a number of countries, states and communities have implemented *Getting to Zero*, *Fast Track Cities*, and *Ending AIDS strategies* with a few being close or achieving the elimination targets well before 2020^38,52-55^. Some are using an active case management approaches that includes social support and immediate treatment with the objective of keeping people healthy and preventing ongoing transmission^38,52-55^. Active case management also includes offering HIV testing for close contacts which, when combined with the offer of treatment and other benefits, has been shown to have higher yields than many other forms of HIV testing. Addressing stigma, discrimination, behavior change (e.g., risk avoidance and delaying age of sexual debut) and other socioeconomic issues (e.g., poverty, access to education) also have a role to play in Ending AIDS. Successful Ending AIDS strategies include a long-term commitment to providing treatment along with a shift in program strategy towards active case management and outbreak control as a bridge to an eventual vaccine or cure.

## The prevention vs. treatment debate: false dichotomy?

*Ending AIDS*, while feasible in most settings, will be difficult and will require strong political leadership and sustained efficient use of limited financial resources. Spending on social priorities such as HIV and health care is often restricted by austerity measures that restrict spending in the public sector, or dwarfed by other priorities, such as military spending. In the current economic and political context, expanding the overall financial envelope for the HIV response is unlikely to happen. The flat-lined, and in many cases decreasing, budgets over the past five years have prompted major donors such as PEPFAR to focus on doing “the right things, in the right places, right now” with an emphasis of maximizing efficient use of existing resources^36^. Debates around resource needs and allocation have traditionally pitted earlier testing and treatment against other prevention interventions^56,57^. The “prevention first” positions are perhaps best exemplified by the traditional mantra that ‘there is no silver bullet”^58^ and “we cannot treat ourselves out of the epidemic”^59^. The recent UNAIDS call for “25% [of spending] for prevention” does not include treatment as a prevention intervention and appears to be the latest effort to carry forward these traditional arguments^60^. Other prevention interventions such as behavior change, pre-exposure prophylaxis (PrEP), condoms, voluntary male circumcision (VMMC), opioid substitution therapy (OST) and needle and syringe programs (NSP) will be necessary, but not sufficient on their own, to end AIDS in many settings^61-63, 66^. The “zero sum game” prevention vs. treatment approach that excludes treatment as prevention of HIV and TB, is contradicted by the science that shows that treatment has been shown to be the most effective prevention intervention. Given its primary therapeutic benefit, it is illogical to exclude treatment from prevention strategies^67^ and ending AIDS will require all of the scientifically proven prevention interventions, including treatment, to be used efficiently. Ending AIDS will require the prioritization and achievement of universal access to early diagnosis and treatment with other prevention interventions such as PrEP, VMMC, condoms, OST and NSP layered on as needed depending on the context.

## Economics of ending AIDS

Although the *90-90-90* targets have been widely accepted and adopted by PEPFAR, The Global Fund, Fast-Track Cities, and others^36,37,38^ there is still controversy surrounding their feasibility, cost and epidemiological impact ^68,69^. These questions have vital importance given the relative flat-lining of current global international and domestic resources and budgeting for HIV at around US$20 billion per annum^70^. The global and national efforts to collect data regarding the resource needs have been extensive but the available information regarding trends, service delivery costs, return on investment, and the relative contribution of international and domestic funding is sparse. Projecting future resource needs for national and global strategic plans vary in their quality and in their framing of the role of treatment and other interventions^23,26,39,71-73^.

UNAIDS-supported national business cases and global needs estimates have restricted access to treatment to those who are severely immunocompromised (e.g., CD4 cell counts below 200, 350 or 500/μL) and commit a majority of resources to ill-defined cost categories such as health systems strengthening, prisons, and social enablers—important categories but whose impact on the epidemic is uncertain and likely to be smaller than other interventions^72^. While prioritizing resources for non-mutually exclusive categories such as social enablers, prison, men who have sex with men is important, however, by doing this the resources for HIV diagnosis and treatment are limited to less than 50% of spending on HIV in many settings^74^. Investment cases need to prioritize HIV testing and immediate access to treatment along with other adjunct prevention options as both a fundamental human right to health and to achieve *90-90-90* by 2020 and *95-95-95* by 2030^26,43,32,74,75,76^.

Reaching the global target of *90-90-90* by 2020 will require a revision of business as usual with resource implications. Current global estimates suggest that around US$20 billion is being invested annually and that this has remained about the same for the past five years^77^. Many of the worst affected countries cannot afford to make large increases in their domestic contributions and continuing support from the international community is still needed. Previous UNAIDS global resource needs projections have ranged from US$32Bn to US$26Bn annually, however, they have limited resources for treatment and include vague non-mutually exclusive budget categories such as enablers and synergies^39,72^. Rough back-of-the-envelope calculations using generous cost per person on ART per year of $300 would result in an 11 billion per annum price tag for the 37 million people living with HIV in the world—far below the UNAIDS 26 billion or 20 billion dollar per annum resource needs estimates^39^. Additionally, this price tag does not allow for discounting, falling drug prices, community based support, time needed to expand the program, and declining incidence. While the true cost of achieving *90-90-90* is unknown, it is likely that resources, time and targets will stay relatively fixed and needs projections for national plans should focus on exploring a more efficient, focused, and evidence-based service delivery model. New costing and epidemiological models need to embrace *90-90-90* as an important milestone towards universal coverage, prioritize testing and treatment, consider the added impact and cost-benefit, if any, of PrEP and other prevention interventions in the context of reaching *90-90-90* and *95-95-95* as essential to the HIV response. Real time programme and population surveillance data should allow for more accurate models and course corrections as needed to ensure the optimal return on investment. Considering that the four largest central banks around the globe, including the United States, the United Kingdom and Japan, printed over $9 trillion in new money in the wake of the 2008 finance crises, it should possible to garner the relatively trivial resources needed to end this humanitarian crisis78.

## Major innovations accelerating the end of AIDS (or the “The Big Five”)

Five major innovations are likely to further accelerate progress of *90-90-90* and the *End of AIDS* (see Figure 2).

**Figure 2:**
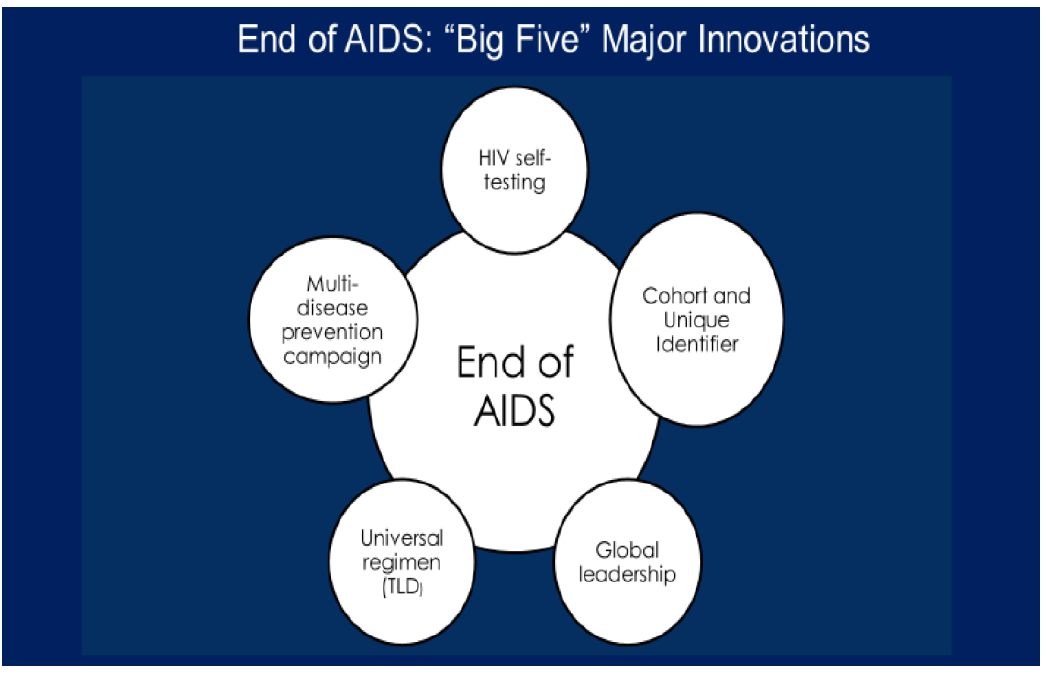
End of AIDS: “Big Five” Major Innovations

## HIV self-tests

Firstly, the recent development of reliable, easy-to-use, HIV self-tests that can be done by people at risk of being infected represents a major advance^50,79^. The new HIV self-tests are highly accurate, rapid, can be done privately without assistance, and in some cases can be read within one minute^80^. Similar to the advent of pregnancy self-tests nearly 40 years ago, this new technology promises to ‘democratize’ access to HIV testing—providing access to testing for people who prefer to do the test in privacy rather than in the traditional health service setting. Building on existing efforts to expand and better target HIV testing delivered by traditional health care workers, access to self-testing will accelerate achievement of the first *90* as people access on-demand HIV testing, seek confirmation to access either HIV treatment or prevention options. Ensuring that people who self-test are welcomed so that their results can be confirmed using a standard HIV diagnosis algorithm is essential. People who are uninfected will need to have access to prevention options and those who are HIV infected can be offered treatment, preferably on the same day^50,51^.

## Universal treatment regimen

The second major innovation is the development of a universal treatment regimen for first-line and as a possible second line option. The introduction of integrase inhibitors and evidence supporting the immediate offer of treatment will have significant impact on lowering barriers to access—this new class of antiretrovirals have improved tolerability with fewer side effects, durability with almost zero resistance, and the potential to be used in two-drug regimens. Although questions regarding the safety of Dolutegravir (DTG) during the early weeks of conception have been raised from a single study in Botswana, more data will be needed to fully evaluate the significance of this *danger signal*. Most researchers have been focused on the potential downsides of being on DTG during conception, only a few have raised questions regarding the downsides of being on an alternate inferior regimen that could lead to less viral suppression, more HIV related morbidity and mortality, and increased HIV transmission to the fetus and partners. Calculations regarding the potential risk of neural tube defect are easier than those concerning use of an inferior regimen but the risk vs benefits should become clearer over the near term. The jury is out, however, it is likely that the TLD regimen will prove to be safe in men and women, including women who are may become pregnant. As with earlier antiretrovirals, prices for integrase inhibitors will likely rapidly decrease and soon they will form the foundation for regimens for middle and lower income countries offering same-day treatment after HIV diagnosis. Announcements regarding price ceilings near $75 dollars for an annual combination regimen of Tenofovir, Lamivudine and Dolutegravir serve as a price guarantee anchor over the near term. However, it also suggests an attempt to guarantee profits given the possibility of market forces leading to far lower prices over the medium and long-term as additional manufacturers join production and nearly all countries adopt it as first and an option for second-line treatment. While some questions have been raised, starting treatment immediately or as soon as possible will likely become the new standard of care^81-84^. This major innovation in treatment practice, particularly when combined with new integrase inhibitor regimens, will provide a sense of urgency to start treatment and render delays for CD4 cell counts or other laboratory tests obsolete. This shift to a “how can we treat people as soon as possible” framing represents a significant shift from the previous WHO “test and wait until immune compromised public health strategy” that led to significant delays and loss to follow-up.

## Comprehensive community-based service delivery model

The third major innovation is the development of a comprehensive integrated community HIV service-delivery model. Studies over the past 10 years have demonstrated that delivering HIV and other health services is feasible in the home or at a community level with very high levels of access. Examples include studies in Western Kenya that offered HIV testing, insecticide treated nets and water filters for over 40,000 people—during a seven day campaign over 80% of the eligible population received services with over a 90% uptake of HIV testing including over 20,000 men^28,30^. More recently the SEARCH project in Uganda and Kenya has implemented an integrated health approach for 117, 711 people in multiple communities that has resulted in 89% of people accepting HIV testing and of these 11 964 (10%) were HIV positive, 4202 (35%) were on ART with 3427 (82%) of those on ART being virally suppressed^84-86^. Additionally, the same group has shown that village leaders, when given minimal support, are able to initiative and lead the delivery of successful community health delivery^85^. Bringing efficient, inexpensive, comprehensive community level health services to people where they live has the potential to ensure access to HIV diagnosis and sustained successful treatment. These three innovations, along with breakthroughs in cloud-based soft-ware for monitoring and evaluation, will improving access to testing and treatment and increase the likelihood of achieving *90-90-90* and *95-95-95* and the *End of AIDS*.

## Standardized M and E that uses a cohort approach combined with unique identifiers

The fourth innovation is the adoption of standardized monitoring and evaluation approaches for expanding access to testing and treatment. Specifically, the clear advantages of using a national cohort approach linked with unique identifiers will allow HIV programs to ensure that everyone who is diagnosed with HIV will have access to comprehensive social services and treatment. This innovation, by allowing movement between clinics to be tracked, will be pivotal to establishing a case management approach to HIV control including social services for those diagnosed with HIV, testing for partners and family, offer of immediate treatment, and testing in places where there is a risk of transmission. Most countries rely on cross-sectional counts of people on treatment at the end of the year and/or cumulative treatment numbers. However, some advanced countries and cities use a national unique identifier and cohort approach that allows the program to follow individuals diagnosed with HIV through the care continuum to viral suppression. Cohort and unique identifiers allow for program accountability for each person with HIV and provide the ability to derive outcomes including healthy and virally suppressed, ill with HIV and non-HIV associated disease, or deaths even if the person transfers care to another clinic. Having relatively precise outcomes should allow for improved stewardship of HIV resources with the ability to shift resources to match burden and to directly link investments with clinical and public health outcomes. In other words, resources can be devoted to supporting people most in need to help them access and stay on treatment. In the near future, block chain technology will allow for an open public ledger for Monitoring and Evaluation, supply chain and other critical information. This would ensure accountability and transparency so that both health authorities and the community could track investments, service delivery and progress towards ending AIDS.

## Unification of global leadership around 95-95-95 target and ending AIDS

The fifth innovation is the potential unification of the global HIV leadership around common disease control objectives, interventions and targets including 95-95-95. PEPFAR has been outstanding in its efforts to focus efforts on expanding access to treatment. National governments have adopted the 90-90-90 targets and many countries are making considerable progress towards universal access. Global leadership around commonly shared definitions of ending AIDS, epidemic control and HIV elimination along with consensus regarding a program strategy that layers other prevention interventions such as PrEP and VMMC onto universal treatment access and viral suppression could allow ending HIV as a serious public health threat in many settings. Like polio, guinea worm, or smallpox eradication, having consensus around epidemic control strategy and objectives, prioritization of resource allocations, and a standardized process for certifying and re-certifying epidemic control will be critical to make epidemic control feasible in many settings. This innovation in leadership will be necessary to move beyond the 50/50/30 challenge—50% of people on ART, 50% of resources for testing and treatment and 30% of people living with HIV virally suppressed.

## Last mile strategies, certification and re-certification

Reaching 90-90-90 and 95-95-95 will transform the epidemiology in many settings leading to a shift in HIV control strategy. Last mile efforts already include active case management approaches reminiscent of traditional infectious disease outbreak control (see Figure 3). Efforts to find the remaining few undiagnosed people living with HIV will require significant innovation. Once diagnosed, programs using last mile strategies rely on a comprehensive social and biomedical response including adherence support, community support, social services, immediate offer of treatment, other prevention options (e.g., condoms), TB prevention and care, and, with the help of the newly person diagnosed, follow-up of other people that may also be infected. Offer of HIV self-tests for the individual’s partners and community-based testing in areas where there is ongoing transmission can be used to find additional people who need access to treatment. When done in a positive fashion with attention to avoid bias, stigma and discrimination, the last mile programs serve to incentivize people to learn their HIV status earlier to access treatment and other services to protect themselves, their families and partners.

**Figure 3:**
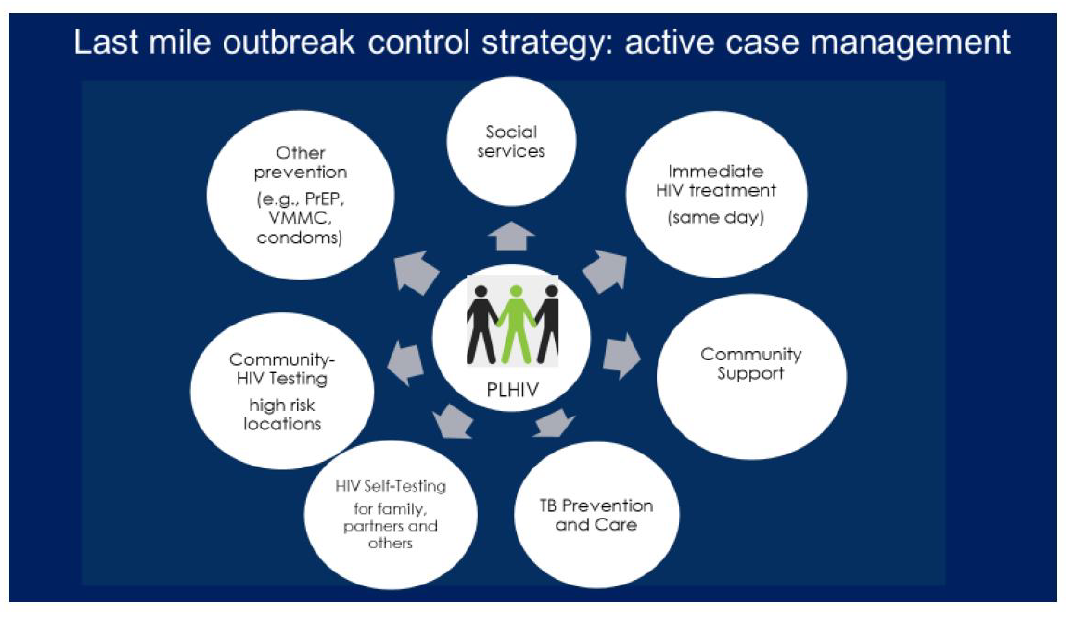
Last mile outbreak control strategy: active case management

Once the epidemic control objectives, definitions and targets are internationally and nationally established it will be important to implement an international certification process to verify achievement of 90-90-90, AIDS incidence, HIV-associated TB incidence (part of AIDS), HIV and non-HIV associated deaths incidence, HIV incidence and prevalence. Although there is already considerable pressure to declare that 90-90-90 and other targets have been achieved, it will be critical that reported denominators and numerators be validated using standardized rigorous criteria^43^. Periodic re-certification will also be necessary since people may be on treatment for decades until a cure or vaccine is discovered and delivered. Lessons for this certification process can be drawn from other disease control efforts including guinea worm eradications, polio eradication, Ebola outbreak mitigation and malaria control among others.

## Conclusion

Global public health security and human rights demand that we focus on ending AIDS for the millions of people living with HIV and impacted by the pandemic. The global response to HIV has been unprecedented and given recent advances in the science and our understanding of the importance of human rights it is feasible to *End AIDS* while supporting people on treatment over the long term. Proof of concept can be seen in many settings including recent population-based surveillance studies that found that HIV prevalence has declined by 34% in Malawi, 31% in Zimbabwe, and 21% Zambia with decreases in incidence as well^87^. *Ending AIDS* as a public health threat includes achieving the *90-90-90* target to ensure successful life-saving treatment for millions of people to both stay healthy and stop HIV transmission. Expanded access to HIV self-testing, life-long treatment with integrase inhibitor based regimens, community delivery of comprehensive HIV services, unified global HIV leadership and increased efficient use of scarce resources, improved monitoring and evaluation and meaningful community engagement and participation will all be necessary to end AIDS.

## Acknowledgements

We would like to thank Jose Zuniga for helpful comments on earlier drafts of this manuscript.

